# Chronic obstructive pulmonary disease and cigarette smoke exposure lead to dysregulated MAIT cell activation by bronchial epithelial cells

**DOI:** 10.1101/2022.02.28.482383

**Authors:** Megan E. Huber, Emily Larson, Taylor N. Lust, Chelsea M. Heisler, Melanie J. Harriff

## Abstract

Chronic obstructive pulmonary disease (COPD) is associated with airway inflammation, increased infiltration by CD8^+^ T lymphocytes, and infection-driven exacerbations. COPD is most commonly caused by cigarette smoke (CS), however the mechanisms driving development of COPD in some smokers but not others are incompletely understood. Lung-resident mucosal-associated invariant T (MAIT) cells play a role in both microbial infections and inflammatory diseases. MAIT cell frequency is reduced in the peripheral blood of individuals with COPD, however the role of MAIT cells in COPD pathology is unknown. Here, we examined MAIT cell activation in response to CS-exposed primary human bronchial epithelial cells (BEC) from healthy, COPD, or smoker donors. We observed significantly higher MAIT cell responses to COPD BEC than healthy BEC. However, COPD BEC stimulated a smaller fold-increase in MAIT cell response despite increased microbial infection. For all donor groups, CS-exposed BEC elicited reduced MAIT cell responses; conversely, CS exposure increased ligand-mediated MR1 surface translocation in healthy and COPD BEC. Our data demonstrate MAIT cell activation is dysregulated in the context of CS and COPD. MAIT cells could contribute to CS- and COPD-associated inflammation through both inappropriate activation and reduced early recognition of bacterial infection, contributing to microbial persistence and COPD exacerbations.

## Introduction

Despite continued smoking cessation programs, smoking remains a major health concern, with eight million deaths in 2017 attributed to tobacco usage^1^. Cigarette smoking is associated with a variety of immunological impacts, including significantly higher susceptibility to microbial infections^2-4^. The components of cigarette smoke act as both pro-inflammatory and immunosuppressive factors that modulate innate and adaptive immunity^3^. For example, cigarette smoke activates caspase 1 to secrete IL-1β and IL-18 *in-vivo*^5-7^, resulting in emphysema and small airway remodeling^8,9^ and accumulation of CD4^+^ and CD8^+^ T cells through IFN-γ signaling^9^. In the context of infection, cigarette smoke inhibits production of pro-inflammatory cytokines in response to microbial infection or LPS stimulation^10^, increases adhesion of *Streptococcus pneumoniae* to bronchial epithelial cells^11^, and delays clearance of *P. aeruginosa*^12^. Conversely, others have observed that repeated cigarette smoke exposure in mice with persistent *S. pneumoniae* airway infection resulted in increased release of pro-inflammatory cytokines including IL-12 and IL-1β, greater bacterial load, and reduced lung function^13^, suggesting that the interplay between cigarette smoke, the airway, and microbial infections is complex.

Cigarette smoking also results in long-term airway changes, evidenced by its role as the primary risk factor for the development of chronic obstructive pulmonary disease (COPD)^14,15^, which itself is the third leading cause of death worldwide^16^. COPD is manifested in a number of clinical phenotypes including small airway disease (e.g. bronchitis) and emphysema, all of which are characterized by chronic inflammation and airflow limitation in the lung and airway^14^. Further complicating COPD pathology are exacerbations, triggered by bacterial or viral colonization and infection, which can increase inflammation and play an important role in the morbidity and mortality associated with COPD^14,15^.

The immune mechanisms underlying the development of airway damage and inflammation leading to COPD in some smokers but not others are poorly defined. Central to this, our understanding of the complex interactions between many cell types is incomplete^17^. CD8^+^ T cells, which are often increased in the lungs of patients with bacterial infections, are the main subset of inflammatory cell increased in the lungs of smokers with COPD compared to asymptomatic smokers^18^. Increased frequencies of CD8^+^ T cells were also observed at the onset of acute exacerbations^19^. Interestingly, chronic cigarette smoke exposure alone resulted in persistent clonal expansion of CD8^+^ T cells in mice^20^. In human COPD lung tissue, CD8^+^ T lymphocytes have increased expression of chemokine receptors, cytotoxic effector molecules, and pro-inflammatory cytokines (reviewed in ^19^). Despite mounting evidence that CD8^+^ T cells are specifically correlated with COPD pathology, the mechanisms underlying the role of CD8^+^ T cells in cigarette smoke- and COPD-mediated pathology remain unclear. Mucosal-associated invariant T (MAIT) cells are an innate-like subset of T lymphocytes that make up a relatively large proportion of the total CD8^+^ T cell population in the blood and lungs in healthy individuals^21^. Interestingly, despite the overall increase in CD8^+^ T cells in COPD, the frequency of both peripheral blood and lung-resident MAIT cells in individuals with COPD is decreased^22-25^. This observation is different from many other infectious and inflammatory lung conditions, and the mechanisms underlying MAIT cell loss in COPD lungs are not yet defined. In fact, little is known about the role of MAIT cells in cigarette smoke- and COPD-associated inflammatory processes.

The antigens presented to MAIT cells by the MHC class I related molecule, MR1, are primarily small molecule metabolites generated during riboflavin biosynthesis by many microbial organisms^26-28^, including those implicated in COPD-associated exacerbations^22,29,30^. MAIT cells can also be activated through both antigen-independent, cytokine-mediated mechanisms^31^. IL-12 and IL-18, the cytokines that elicit this type of antigen-independent response, are among those produced by airway epithelial cells and other inflammatory cells in the context of cigarette smoke and COPD^5,7,32^. The direct impact of cigarette smoking and COPD on MR1 antigen presentation and subsequent MAIT cell responses to infected airway cells is unknown, however. We hypothesized that exposure of bronchial epithelial cells (BEC) to cigarette smoke and the inflammatory COPD airway environment would result in dysregulated MAIT cell responses through altered MR1 function, contributing to inflammation and exacerbation. We found that exposure of BEC to cigarette smoke decreased both microbe-independent and microbe-dependent responses. Furthermore, BEC from COPD lungs induced greater MAIT cell responses compared to healthy controls. Exposure to cigarette smoke did not affect transcriptional expression of *MR1*, but did result in increased MR1 surface expression, suggesting that smoking may interfere with the ability of MR1 to encounter microbial ligands. Our data demonstrate that impaired interactions between airway epithelial cells and MAIT cells, resulting in dysregulated release of pro-inflammatory cytokines and other molecules, may play a role in COPD-associated inflammation in the context of both cigarette smoke as well as bacterial colonization and infection.

## Results

### BEC from COPD lungs induce increased microbe-independent, MR1-dependent activation of MAIT cells

Inappropriate MR1 antigen presentation and activation of lung-resident MAIT cells could contribute to the inflammatory airway environment present in COPD airways and following cigarette smoking. As such, we tested the ability of MAIT cells to respond to primary human BEC from the lungs of COPD or smoker donors compared to healthy controls, and in the context of cigarette smoke exposure. BEC were isolated from the lungs of healthy (N=7), COPD (N=6), or smoker (N=6) donors between the ages of 41 and 73 (Table 1). BEC from these donors were incubated with a previously described MAIT cell clone (D426 G11)^33,34^ following treatment with cigarette smoke extract (CSE) and infection with *Mycobacterium smegmatis* or *Streptococcus pneumoniae* in an ELISPOT assay with IFN-γ production by the MAIT cell clone as the readout. A linear mixed effects model with square root transformation of the IFN-γ spot forming units (SFU) was used to analyze the data for significant effects of donor BEC groups on MAIT cell responses.

**Table 1.**
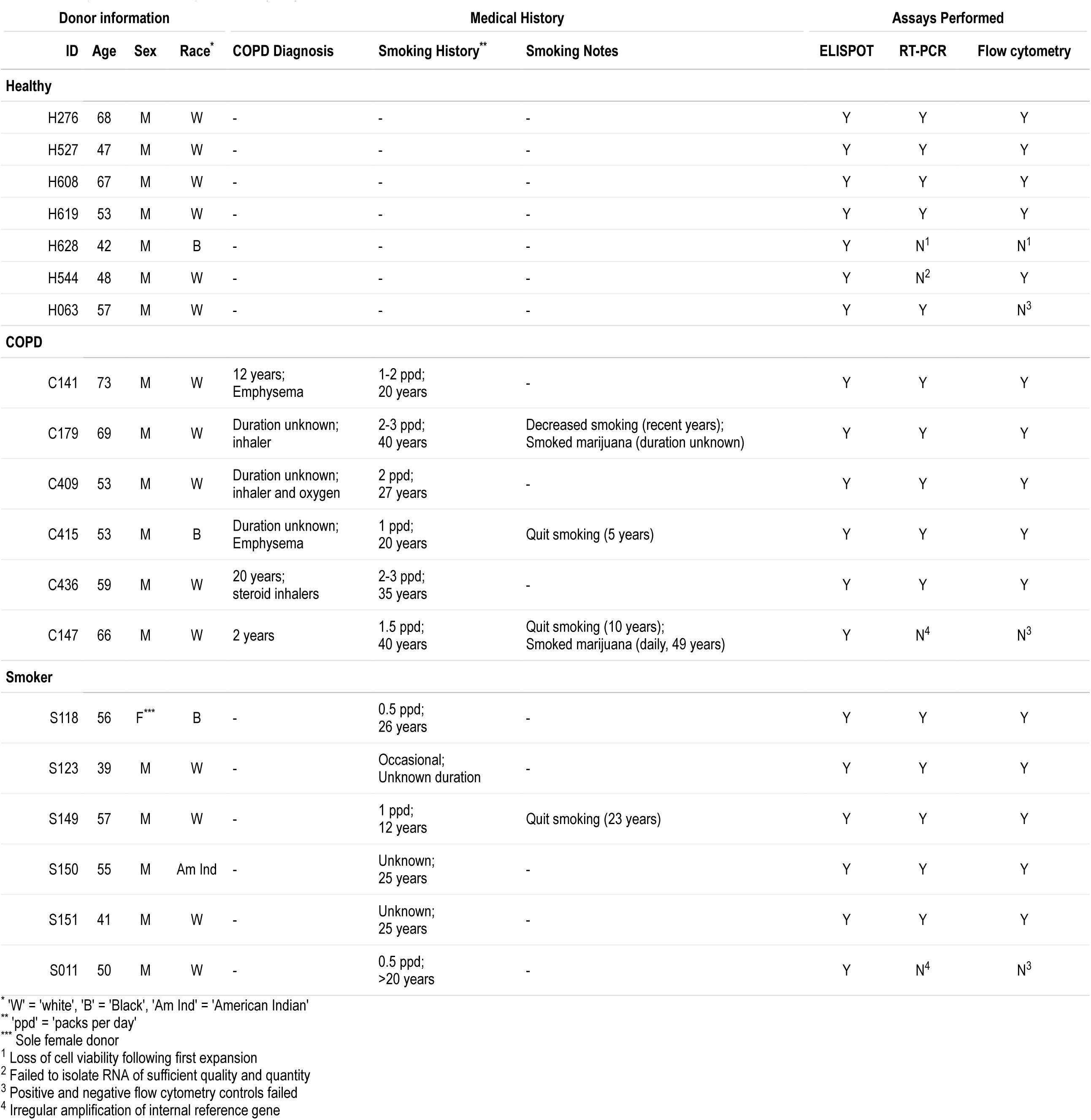
Description of bronchial epithelial cell (BEC) donors

We first analyzed the response of the MAIT cell clone to uninfected BEC. We observed significantly greater microbe-independent IFN-γ SFU in response to BEC from COPD donors than healthy or smoker donors (p=0.0416, Figure 1a, Table 2). These microbe-independent MAIT cell responses to COPD donors were greater than responses observed to the uninfected bronchial epithelial cell line (BEAS-2B) control (Figure S1). There were no differences in the MAIT cell response to smoker donors compared to healthy controls (p=0.5173, Figure 1a, Table 2). We hypothesized that an increase in pro-inflammatory cytokines capable of mediating MAIT cell responses such as IL-18^31^, which is produced by primary BEC from the lungs of subjects with COPD^5,6,32^, could induce increased MR1-independent MAIT cell responses absent microbial antigens. To determine whether stimulation of IFN-γ production by the MAIT cells occurred through MR1-or cytokine-dependent pathways, we used antibodies to block MR1 or IL-12 and IL-18 in BEC from a representative healthy and COPD donor. There was almost complete blockade of the IFN-γ SFU response for both the healthy and COPD donors in the presence of the 26.5 α-MR1 antibody, with very little impact of blocking IL-12 and IL-18 (Figure 1b-c). This suggests that despite the lack of antigen from microbial infection, there are nonetheless MR1-dependent MAIT cell responses to primary BEC from all donors. Further, these MR1-dependent responses are increased in the context of cells from COPD lungs.

**Table 2.** Statistical analysis of ELISPOT data: Fixed effects results from linear mixed model

| Variable | p value |
| --- | --- |
| Healthy | (Intercept) |
| COPD | 0.0416 |
| Smoker | 0.5173 |
| CSE | 0.5279 |
| *Msm* | <0.0001 |
| *Sp* | <0.0001 |
| COPD:CSE | 0.0003 |
| Smoker:CSE | <0.0001 |
| COPD:*Msm* | <0.0001 |
| Smoker:*Msm* | 0.8718 |
| COPD:*Sp* | 0.4193 |
| Smoker:*Sp* | 0.2432 |
| CSE:*Msm* | <0.0001 |
| CSE:*Sp* | 0.0482 |

Acronyms: *Msm* = *M. smegmatis* *Sp* = *S. pneumoniae* CSE = cigarette smoke extract

**Figure 1.**
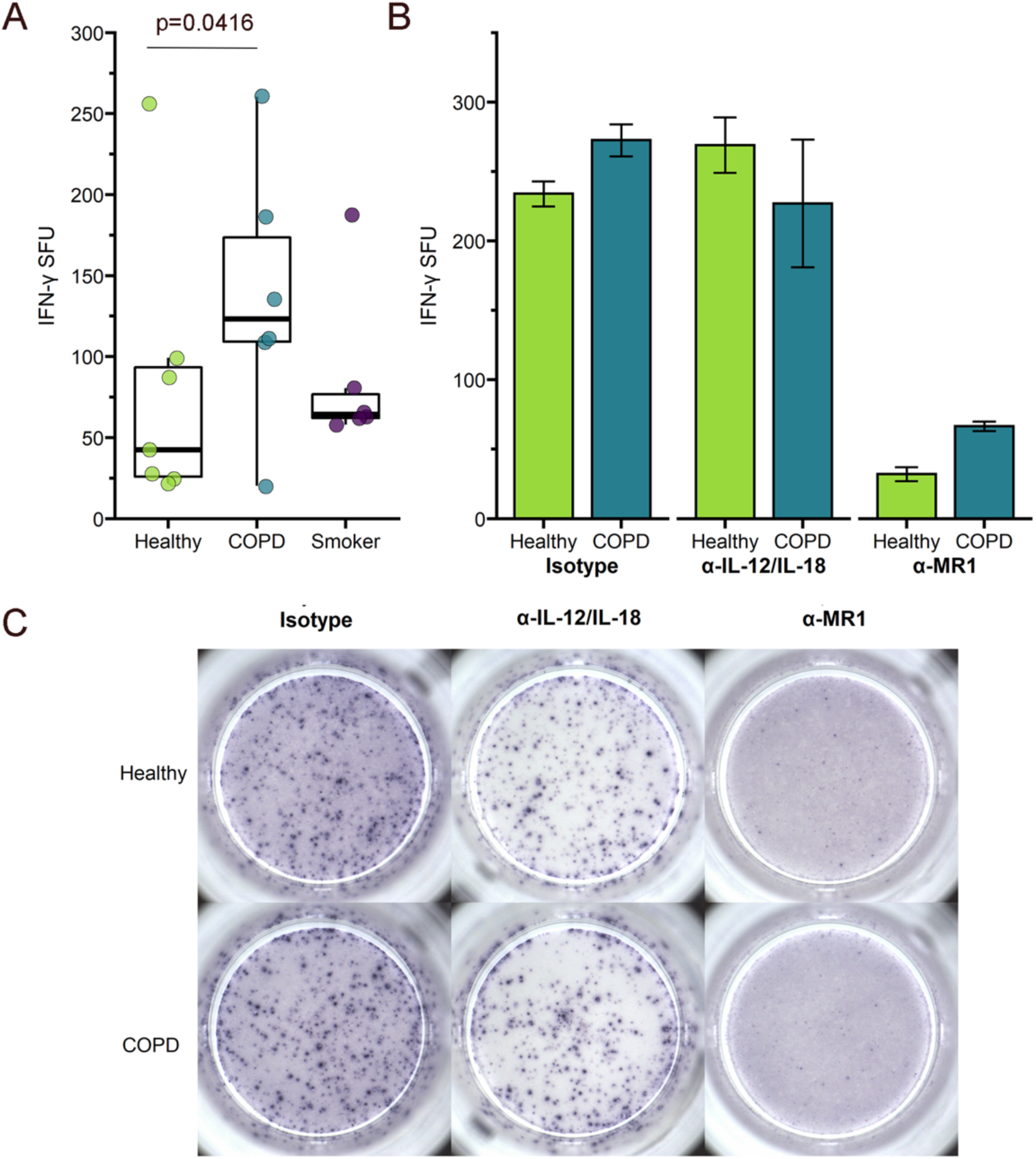
Primary BEC elicit microbe-independent, MR1-dependent responses by MAIT cells. a) Primary BEC from healthy (n=7), COPD (n=6), or smoker (n=6) donors were incubated with the D426 G11 MAIT cell clone in an ELISPOT assay with IFN-γ production as the readout. Data points are the mean IFN-γ spot-forming units (SFU) of two technical replicates per donor. Statistical analysis was performed as described in the experimental procedures and is summarized in Table 2. b-c) BEC from a representative healthy and COPD donor were incubated with blocking antibodies to IL-12/IL-18 or MR1 five hours prior to addition of the MAIT cells in an IFN-γ ELISPOT assay. Results are presented as b) the mean of two experimental replicates and c) representative ELISPOT well images.

We did observe diffuse IFN-γ staining haze in all ELISPOT wells containing both BEC and MAIT cells (Figure 1c). This haze was completely abrogated in the context of IL-12 and IL-18 blocking for both donors, demonstrating that there are likely cytokine-mediated MAIT cell responses to the primary BEC in addition to the MR1-dependent responses. Quantification of non-spot forming IFN-γ is not possible in the context of an ELISPOT assay. Therefore, we were unable to determine whether there was also a meaningful difference in this cytokine-dependent response to the healthy or COPD donor BEC. We did, however, perform an assay to detect IL-18 secretion by a representative healthy, COPD, and smoker donor. All donors produced less than 2pg/mL of IL-18, with no difference between the donors (Figure S2). Taken together with the abrogation of IFN-γ spots in the presence of the α-MR1 antibody, our data suggest that microbe-independent MAIT cell activation is largely mediated through MR1-dependent mechanisms and is increased in response to COPD BEC.

### BEC from COPD lungs induce a decreased fold-change in microbe-dependent, MR1-dependent activation of MAIT cells

We next looked at MAIT cell responses to BEC from these same donors infected with the pneumonia-causing pathogen *S. pneumoniae* (*Sp*), or with *M. smegmatis* (*Msm*) as a positive control. As expected, the linear mixed effects modeling showed that MAIT cell responses to the *Sp*-or *Msm*-infected healthy donor BEC were significantly greater than responses to uninfected BEC (p<0.0001, Figure 2a, Table 3). Similar to the microbe-independent ELISPOT assays, the MAIT cell IFN-γ SFU responses to infected BEC required MR1, as demonstrated by nearly complete blocking in the presence of the 26.5 α-MR1 antibody (Figure 2b). Overall, *Msm-* or *Sp-* infected BEC from COPD donors induced higher, but not statistically significant, MAIT cell responses than infected BEC from healthy or smoker donors (Figure 2a, Table 2). To further explore this observation, we enumerated microbial infection of BEC from healthy, COPD or smoker donors using fluorescence microscopy. We observed significantly more *Sp* cocci associated with BEC from COPD lungs compared to BEC from healthy or smoker lungs (p<0.0001, Figure 3a-b).

**Table 3.**
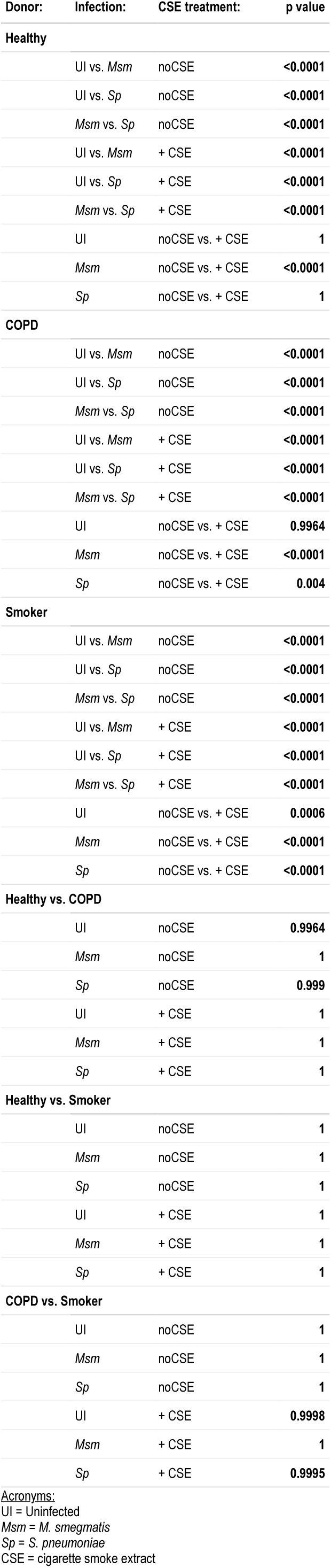
Statistical analysis of ELISPOT data:

| Donor: | Infection: | CSE treatment: | p value |
| --- | --- | --- | --- |
| Healthy |  |  |  |
|  | UI vs. *Msm* | noCSE | <0.0001 |
|  | UI vs. *Sp* | noCSE | <0.0001 |
|  | *Msm* vs. *Sp* | noCSE | <0.0001 |
|  | UI vs. *Msm* | + CSE | <0.0001 |
|  | UI vs. *Sp* | + CSE | <0.0001 |
|  | *Msm* vs. *Sp* | + CSE | <0.0001 |
|  | UI | noCSE vs. + CSE | 1 |
|  | *Msm* | noCSE vs. + CSE | <0.0001 |
|  | *Sp* | noCSE vs. + CSE | 1 |
| COPD |  |  |  |
|  | UI vs. *Msm* | noCSE | <0.0001 |
|  | UI vs. *Sp* | noCSE | <0.0001 |
|  | *Msm* vs. *Sp* | noCSE | <0.0001 |
|  | UI vs. *Msm* | + CSE | <0.0001 |
|  | UI vs. *Sp* | + CSE | <0.0001 |
|  | *Msm* vs. *Sp* | + CSE | <0.0001 |
|  | UI | noCSE vs. + CSE | 0.9964 |
|  | *Msm* | noCSE vs. + CSE | <0.0001 |
|  | *Sp* | noCSE vs. + CSE | 0.004 |
| Smoker |  |  |  |
|  | UI vs. *Msm* | noCSE | <0.0001 |
|  | UI vs. *Sp* | noCSE | <0.0001 |
|  | *Msm* vs. *Sp* | noCSE | <0.0001 |
|  | UI vs. *Msm* | + CSE | <0.0001 |
|  | UI vs. *Sp* | + CSE | <0.0001 |
|  | *Msm* vs. *Sp* | + CSE | <0.0001 |
|  | UI | noCSE vs. + CSE | 0.0006 |
|  | *Msm* | noCSE vs. + CSE | <0.0001 |
|  | *Sp* | noCSE vs. + CSE | <0.0001 |
| Healthy vs. COPD |  |  |  |
|  | UI | noCSE | 0.9964 |
|  | *Msm* | noCSE | 1 |
|  | *Sp* | noCSE | 0.999 |
|  | UI | + CSE | 1 |
|  | *Msm* | + CSE | 1 |
|  | *Sp* | + CSE | 1 |
| Healthy vs. Smoker |  |  |  |
|  | UI | noCSE | 1 |
|  | *Msm* | noCSE | 1 |
|  | *Sp* | noCSE | 1 |
|  | UI | + CSE | 1 |
|  | *Msm* | + CSE | 1 |
|  | *Sp* | + CSE | 1 |
| COPD vs. Smoker |  |  |  |
|  | UI | noCSE | 1 |
|  | *Msm* | noCSE | 1 |
|  | *Sp* | noCSE | 1 |
|  | UI | + CSE | 0.9998 |
|  | *Msm* | + CSE | 1 |
|  | *Sp* | + CSE | 0.9995 |

Acronyms: UI = Uninfected *Msm* = *M. smegmatis* *Sp* = *S. pneumoniae* CSE = cigarette smoke extract

**Figure 2.**
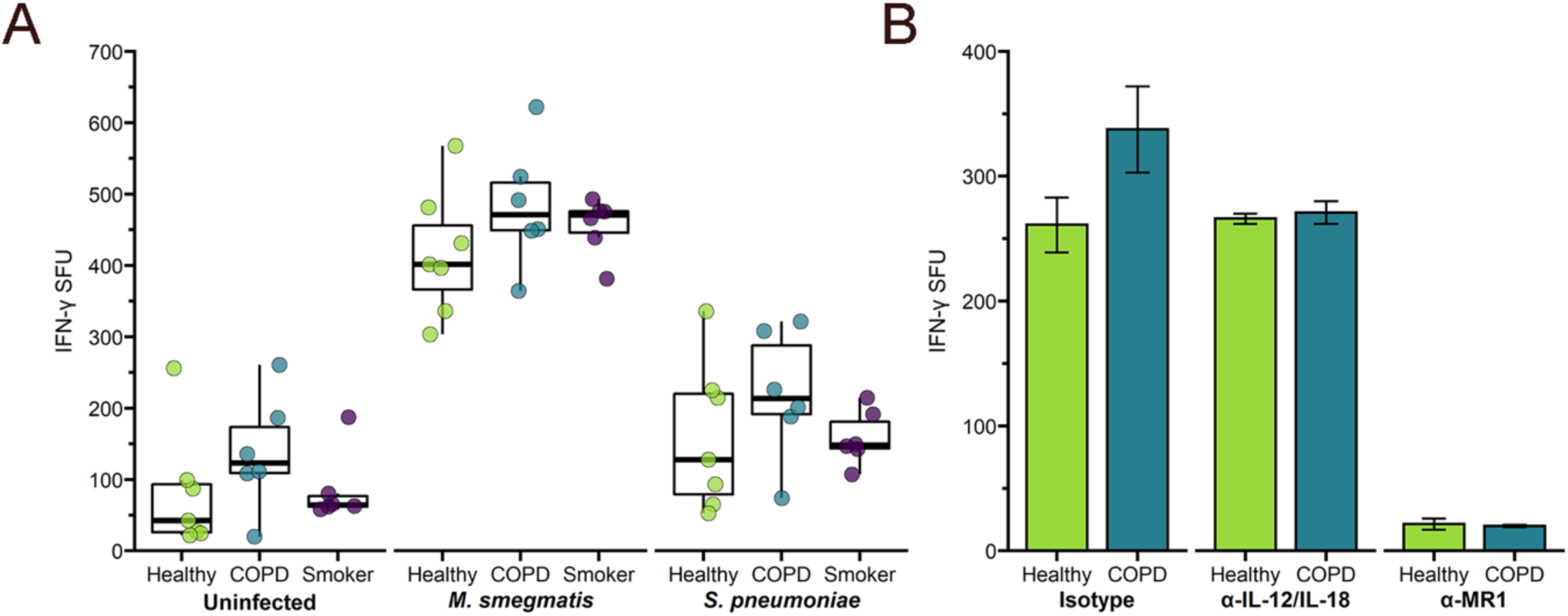
Increased microbe-dependent, MR1-dependent MAIT cell response to infected BEC from COPD donors. a) Primary BEC from healthy, COPD, or smoker donors were infected with media control, *M. smegmatis* (0.1μl/well), or *S. pneumoniae* (MOI 20) for one hour prior to addition of MAIT cells in an IFN-γ ELISPOT assay. Data points are the mean IFN-γ spot-forming units (SFU) of two technical replicates per donor. Statistical analysis was performed as described in the experimental procedures and is summarized in Table 3. b) BEC from a representative healthy and COPD donor were incubated with blocking antibodies to IL-12/IL-18 or MR1 one hour prior to infection with *S. pneumoniae* (MOI 20) and subsequent addition of the MAIT cell clones in an IFN-γ ELISPOT assay. Results are presented as the mean of two experimental replicates.

**Figure 3:**
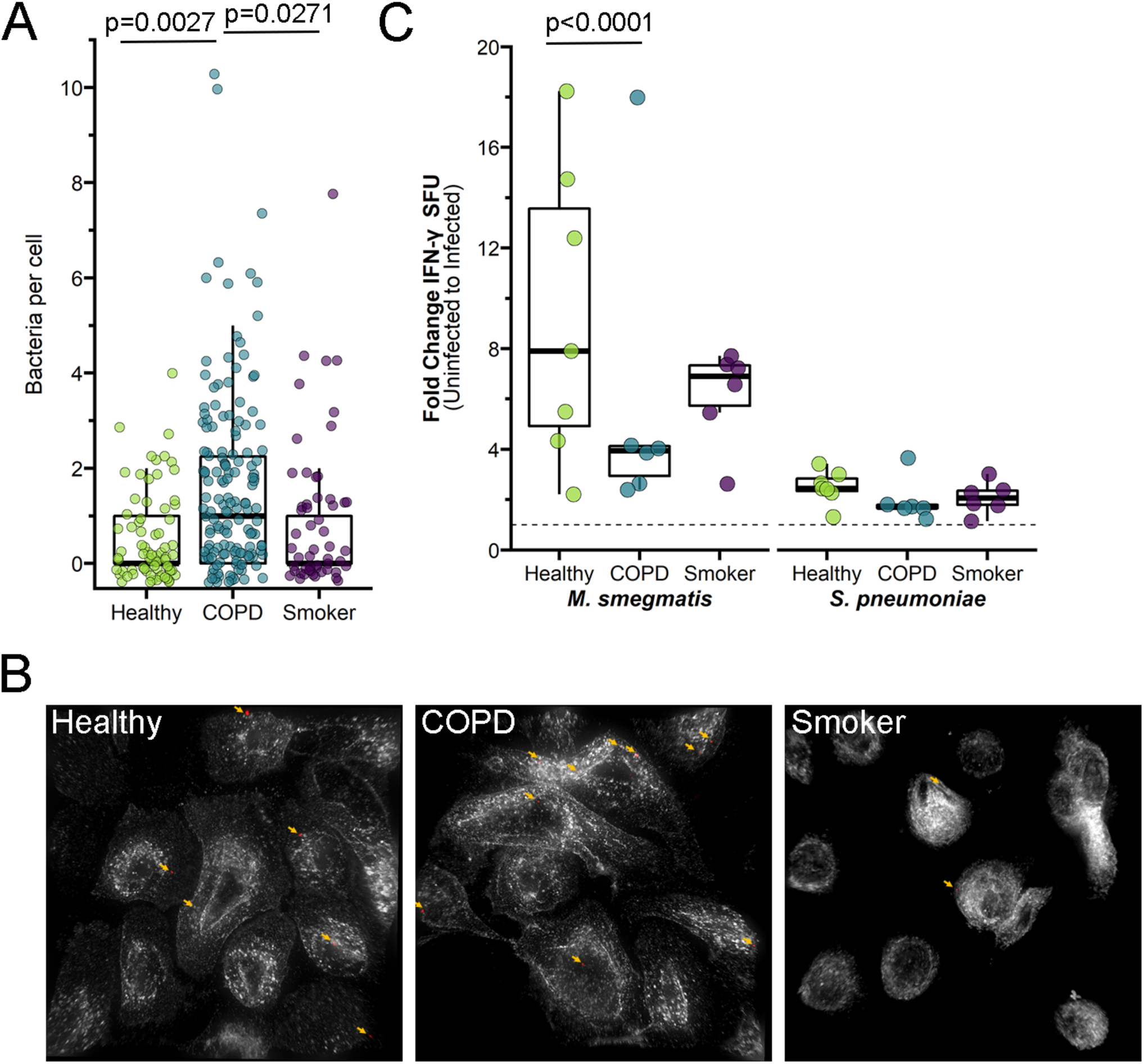
Increased infection of primary BEC from COPD donors. a-b) BEC from representative healthy, COPD, or smoker donors were infected with fluorescently labeled *S. pneumoniae* for three hours. Fixed cells were stained with DAPI and α-MHC-Ia antibody to label the cell surface. Approximately 20 fields per donor were selected without bias based on nuclear stain, and whole cells within these fields were then analyzed by Imaris to enumerate the number of bacteria associated with individual cells. a) Data points indicate individual cells, analyzed by one-way ANOVA statistical analysis. b) Representative images of *S. pneumoniae*-infected primary BEC. White = MHC-Ia surface staining. Red pseudocolor = fluorescent *S. pneumoniae*. Arrows indicate adherent *S. pneumoniae* (yellow) enumerated for analysis. c) IFN-γ SFU fold change between no-treatment control and *M. smegmatis-* or *S. pneumoniae-*infected primary BEC from healthy, COPD, or smoker donors. Raw data shown in Figure 2a. Statistical analysis was performed as described in the experimental procedures and is summarized in Table 2.

To quantify the infection-mediated increase in MAIT cell IFN-γ production and take into account the differences in antigen-independent activation of MAIT cells between the BEC donor groups, we compared the pairwise fold change in IFN-γ SFU responses between uninfected and infected donor BEC (Figure 3c). Surprisingly, the infection-mediated increase in MAIT cell responses to infected COPD donor BEC was reduced in comparison to fold-change responses to healthy and smoker donor BEC. Therefore, despite significantly greater bacterial infection per cell and overall higher induction of MAIT cell IFN-γ production, COPD donor BEC stimulated a weaker MAIT cell response upon infection. These results suggest that MR1 antigen presentation is impaired in infected BEC from COPD lungs.

### Exposure to cigarette smoke decreases MAIT cell activation in response to primary BEC from healthy, COPD, and smoker lungs

We next examined whether treating primary BEC with cigarette smoke impacts microbe-independent MAIT cell responses. BEC were treated with 30% cigarette smoke extract (CSE) prior to infection and subsequent incubation with MAIT cell clones in an IFN-γ ELISPOT assay. For healthy and COPD donor BEC, CSE treatment did not significantly affect overall MAIT cell IFN-γ responses (Figure 4a, Table 3). Interestingly, when comparing the fold change in IFN-γ SFU, BEC from COPD and smoker donors induced significantly lower MAIT cell responses after incubation with CSE (Figure 4b, Table 2). In other words, the magnitude of the CSE-mediated decrease in MAIT cell response was significantly greater for COPD and smoker BEC than healthy donors.

**Figure 4:**
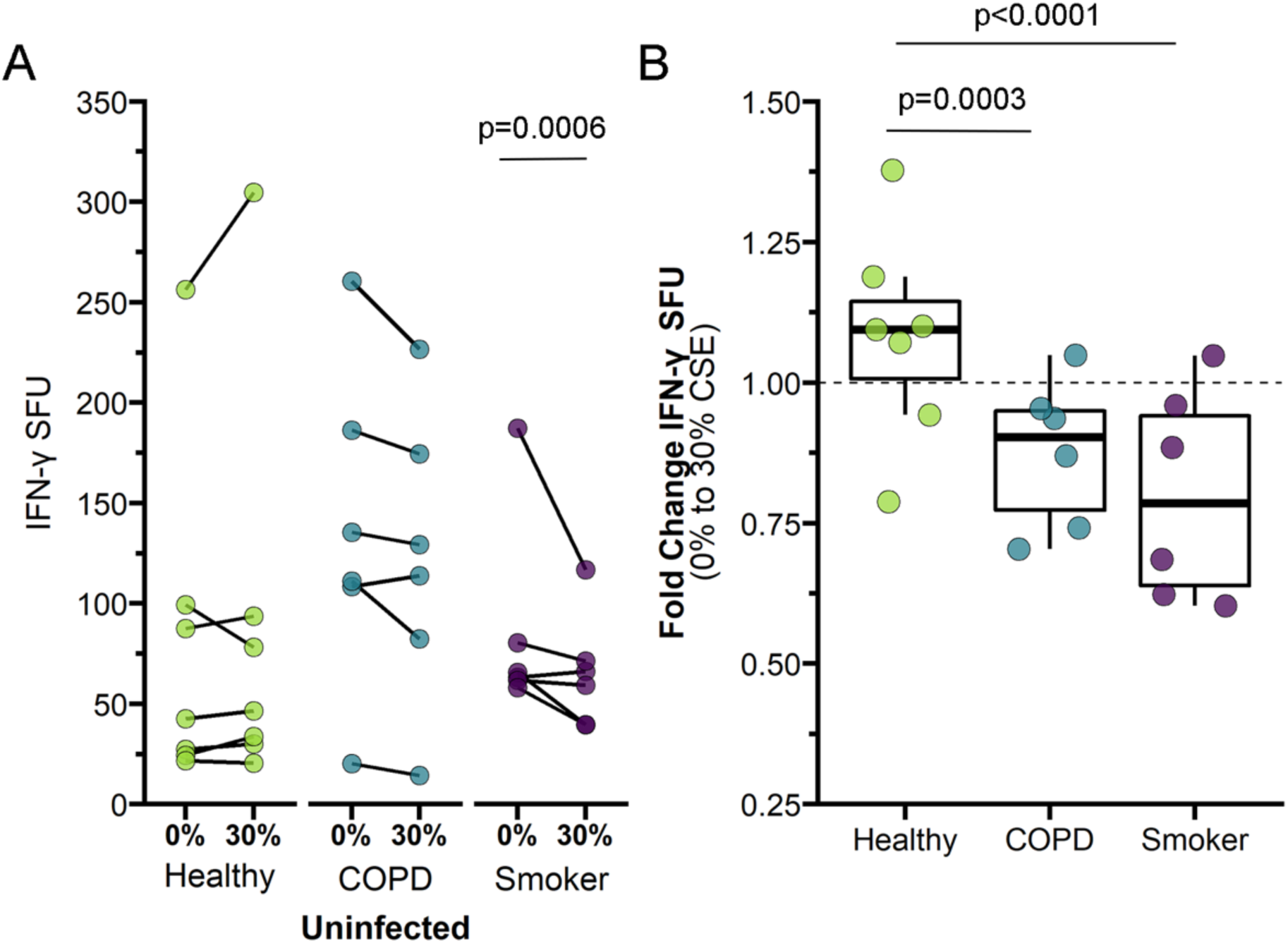
Decreased MAIT cell responses to primary BEC following treatment with cigarette smoke extract (CSE). a-b) Primary BEC from healthy (n=7), COPD (n=6), or smoker (n=6) donors were incubated with media containing 0% or 30% CSE for three hours prior to the addition of MAIT cells in an IFN-γ ELISPOT assay. Statistical analysis was performed as described in the experimental procedures and is summarized in Tables 2-3. a) Data points are the mean IFN-γ spot-forming units (SFU) of two technical replicates, paired by individual donor. b) IFN-γ SFU fold change between 0% CSE- and 30% CSE-treated primary BEC from healthy, COPD, or smoker donors, calculated pairwise by donor.

We next explored the impact of CSE treatment in combination with bacterial infection. Notably, MAIT cell responses to infected BEC from all donor groups were significantly reduced by CSE treatment (Figure 5a-b, Table 2-3). The decreased response with CSE treatment was unexpected, given previous reports indicating that cigarette smoke treatment increases *Sp* infection of respiratory tract epithelial cells^35-37^. As such, we examined the efficiency of *Sp* infection in these cells as described above. CSE treatment increased the number of *Sp* cocci per BEC from healthy (p<0.001) and COPD donor groups (p=0.029, Figure 5d-e). In the context of this increased infection, our observation of decreased IFN-γ SFU response to CSE-treated cells suggested cigarette smoke may downregulate MR1 antigen presentation to MAIT cells. The average number of microbes per BEC was significantly greater for CSE-treated BEC from COPD donors than from healthy (p-0.034) or smoker donors (p=0.0053, Figure 5d-e). There were no significant donor group differences in the fold change IFN-γ response to CSE-treated, infected BEC (Figure 5c), these results suggesting that the combination of infection and CSE treatment may affect healthy BEC similarly to COPD BEC. Interestingly, CSE treatment did not significantly affect *Sp* infection of BEC from smoker donors (Figure 5d-e) despite reduced MAIT cell responses (Figure 5b), suggesting that cigarette smoke alteration of *Sp* infection and downstream MAIT cell responses may occur through varied mechanisms that differ in the context of acute versus chronic smoke exposure. Together, our data suggest a complex role for cigarette smoke in modulating MR1 antigen presentation to MAIT cells.

**Figure 5:**
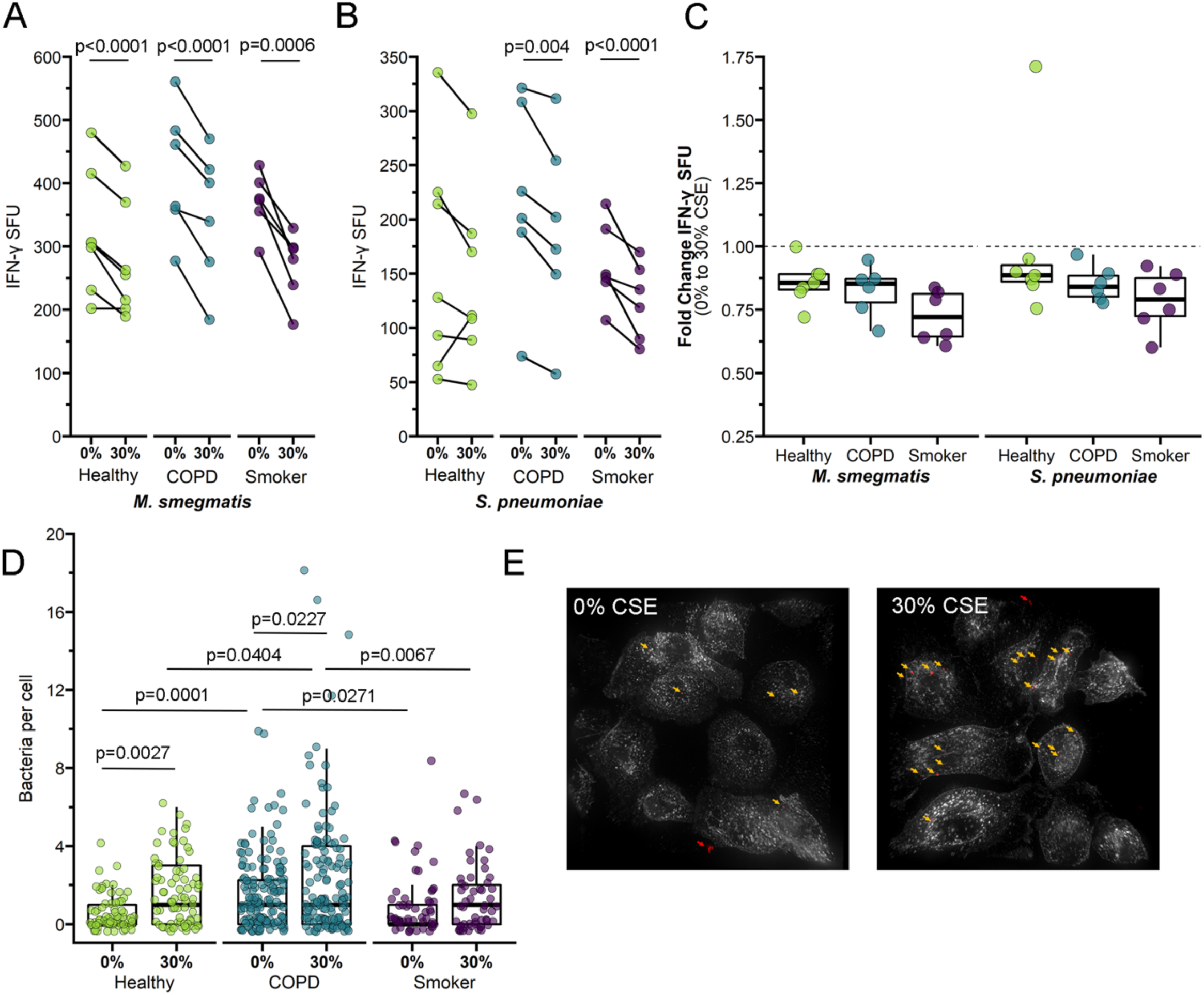
Reduced MAIT cell responses to infected BEC following treatment with CSE. a-b) Primary BEC from healthy (n=7), COPD (n=6), or smoker (n=6) donors were incubated with media containing 0% or 30% CSE for three hours. BEC were infected with *M. smegmatis* (a, 0.05μl/well) or *S. pneumoniae* (b, 20 MOI) for one hour prior to the addition of MAIT cells in an IFN-γ ELISPOT assay. Statistical analysis was performed as described in the experimental procedures and is summarized in Table 2-3. c) Fold change of a) and b) IFN-γ SFU between 0% CSE- and 30% CSE-treated primary BEC infected with *M. smegmatis* or *S. pneumoniae*, calculated pairwise by donor. d-e) BEC from representative healthy, COPD, or smoker donors were incubated with medium containing 0% or 30% CSE for three hours, then infected with fluorescently labeled *S. pneumoniae* for three hours. Fixed cells were stained with DAPI and α-MHC-Ia antibody to label the cell surface. Approximately 20 fields per donor were selected without bias based on nuclear stain, and whole cells within these fields were then analyzed by Imaris to enumerate the number of bacteria associated with individual cells. d) Representative images of *S. pneumoniae*-infected healthy BEC treated with 0% or 30% CSE. White = MHC-Ia surface staining. Red pseudocolor = fluorescent *S. pneumoniae*. Arrows indicate adherent *S. pneumoniae* (yellow) enumerated for analysis or extracellular microbes (red) excluded for analysis. e) Data points indicate individual cells, analyzed by one-way ANOVA statistical analysis. 0% CSE results from Figure 3a are included as reference.

### Acute CS exposure does not impact transcriptional regulation of MR1

Our ELISPOT and infection results suggested that MR1-dependent MAIT cell activation was impacted in cells from the COPD lung environment and following acute treatment with CSE. We considered the possibility that altered MR1 expression in these cells could explain these changes. Although MR1 expression has been confirmed in all cell types studied to-date^38^, nearly all analyses of MR1 expression and regulation have focused on the surface expression of MR1 protein. There are a limited number of studies examining *MR1* gene expression in bulk cells from the lung parenchyma or peripheral blood of COPD donors^24,25^, however we are unaware of any analysis of the impact of COPD or CS exposure on *MR1* expression in primary BEC. Therefore, we looked at *MR1* gene expression in BEC from COPD and smoker lungs compared to healthy controls and sought to determine if exposure to CSE had any impact on *MR1* mRNA expression in BEC from all donors. We isolated mRNA from BEC following CSE treatment and corresponding control conditions, and measured expression of *MR1* and the internal control *HPRT1*. Baseline C_t_ values and ΔC_t_ analysis of MR1 mRNA across all donors revealed significantly higher expression of MR1 mRNA in smoker donors compared to healthy controls at baseline (Figure 6a, p=0.0200). The smaller ΔCt value for COPD BEC compared to healthy donors suggested higher *MR1* expression, however this difference was not significant (Figure 6a). Using the 2^-ΔΔCt^ method, we then determined the fold increase in *MR1* expression within each donor in the context of CSE treatment relative to no treatment (Figure 6b). For all BEC donor groups, paired comparisons demonstrated there were no statistically significant impacts of acute CSE exposure on *MR1* expression. Although there were no significant impacts of these treatments alone when examining paired responses for individual donors, ANOVA analysis of the ΔC_t_ values between the groups as a whole indicated that, similar to the baseline condition, BEC from smoker donors treated with CSE still expressed significantly greater *MR1* compared to healthy donors (p=0.0319, Figure 6b). Taken together, these results do not demonstrate a consistent role for baseline expression or CSE-mediated transcriptional regulation of MR1 in the observed ability of MAIT cells to respond to BEC. While increased *MR1* expression in BEC from COPD lungs could explain the increase in microbe-independent responses to these cells, these findings are not consistent in BEC from smoker lungs.

**Figure 6:**
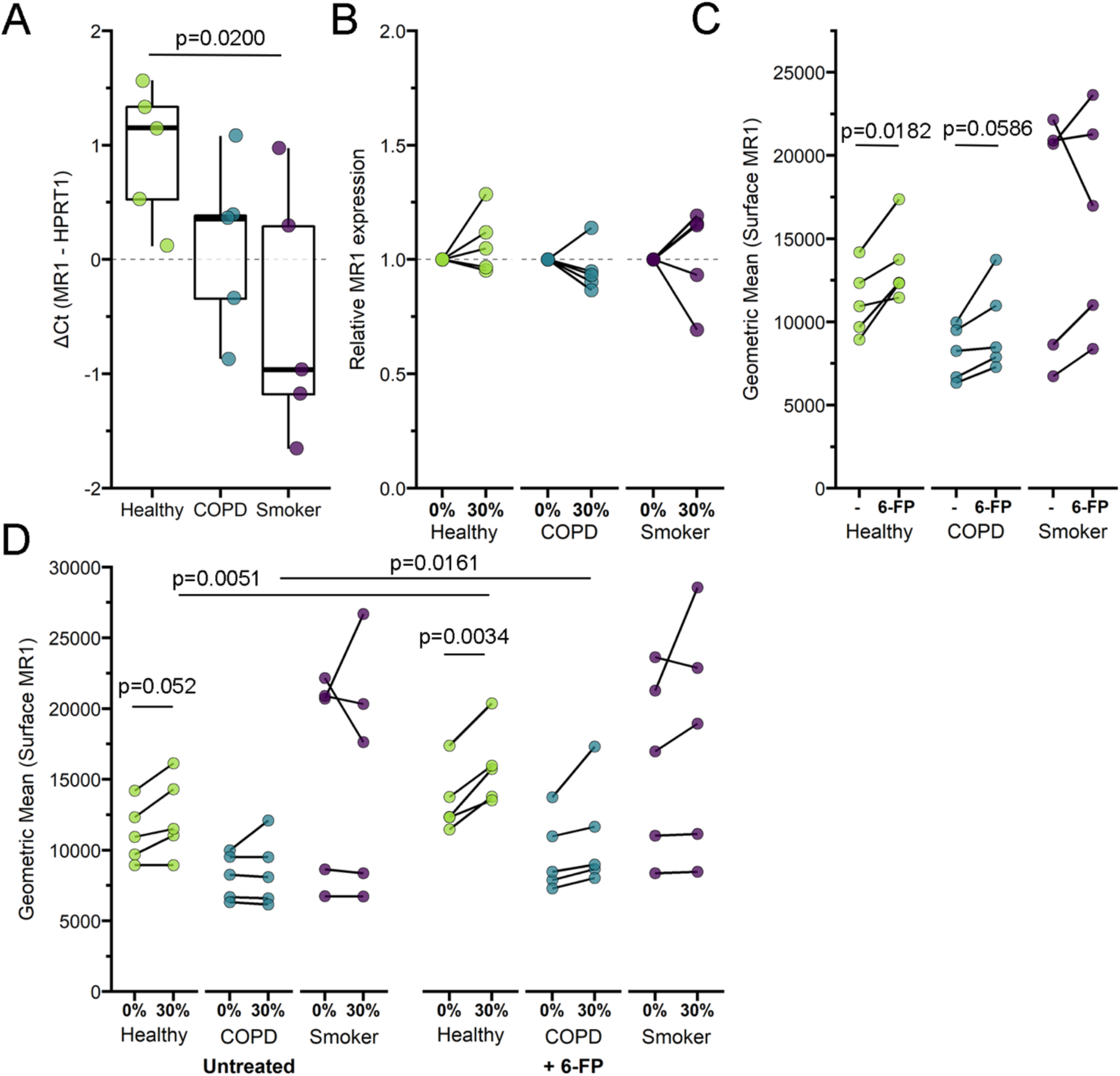
Increased MR1 expression in primary BEC exposed to cigarette smoke. a) RNA was isolated from healthy, COPD, or smoker donor BEC (n=5) and RT-PCR was performed to detect amplification of *MR1* and the internal control, *HPRT1*. Data points are the mean ΔC_t_ of three technical replicates per donor. Statistical analysis performed by two-way ANOVA with Bonferroni’s multiple comparisons. b) Primary BEC from healthy, COPD, or smoker donors (n=5) were incubated with 0% or 30% CSE for 3 hours. RNA was isolated from BEC and RT-PCR was performed to detect amplification of *MR1* and the internal control, *HPRT1*. Relative mRNA 2^-ΔΔCt^ calculations were performed relative to no-treatment pairwise control and *HPRT1* expression. Two tailed paired t-tests were performed to determine statistical significance; p value > 0.05 for all comparisons. c-d) Primary BEC from healthy, COPD, or smoker donors (n=5) were d) incubated with 0% or 30% CSE for three hours, then c-d) incubated overnight with the MAIT cell ligand 6-formylpterin (6-FP) prior to harvest and staining for surface expression of MR1 by flow cytometry. Two-tailed paired t tests were performed to determine statistical significance.

### Acute cigarette smoke exposure increases MR1 surface translocation in BEC

Although the *MR1* mRNA expression was not significantly different, other impacts to intracellular function in the context of CS or COPD could result in changes to surface MR1 protein expression. To examine this possibility, we measured the surface expression of MR1 on primary BEC at basal levels and following induction of MR1 surface translocation through treatment with the ligand 6-formylpterin (6-FP). Consistent with our previous studies, the level of endogenous MR1 surface expression in *ex-vivo* primary BEC is relatively low compared to cell lines^39^, particularly those that overexpress MR1^40^. As such, we included BEAS-2B cells overexpressing MR1 in each assay as a control to confirm detection and surface translocation of MR1 (Figure S3a-b). In our primary BEC, despite the expected low MR1 surface expression, we were nonetheless able to detect 6-FP-mediated increases in surface MR1 in all of our donors (Figure 6c). In healthy BEC, there was a significant increase in surface MR1 following 6-FP treatment (p=0.0182). Although not significant, we also observed a modest increase for each COPD and smoker BEC donor treated with 6-FP. We then assessed the role of acute exposure to CSE in modulating these processes. As with *MR1* mRNA expression, basal expression of surface MR1 in healthy and COPD cells was not impacted by CSE treatment (Figure 6d). However, the 6-FP-mediated increase in surface MR1 was increased in the context of CSE treatment for healthy (p=0.0051) and COPD donors (p=0.0161). Interestingly, we observed no significant pairwise changes from CSE treatment alone or in combination with 6-FP for smoker donor BEC (Figure 6d), further indicating that MR1 expression is differentially affected by acute CSE exposure, long-term cigarette smoking, and intracellular changes induced in BEC during the development of COPD. Together, our results demonstrate that acute exposure to cigarette smoke may impact ligand-dependent surface translocation of MR1.

## Discussion

MAIT cells are an evolutionarily conserved subset of T cells present in high proportions in human blood and peripheral mucosal sites. While MAIT cells were first described for their role in recognizing and responding to microbial infection^33,41^, evidence continues to grow for their role in inflammatory non-infectious diseases^42^. Furthermore, MAIT cells have now been implicated in the homeostasis and repair of various mucosal barrier tissues, including the lung^43^. MAIT cell functions may be relevant both to the cigarette smoke-mediated development of airway inflammation resulting in COPD pathologies and to airway exacerbations common in COPD. Of note, numerous groups have observed decreased MAIT cell frequencies in both the peripheral blood and lungs of individuals with COPD^22,23,25^, which is contrary to the increase in MAIT cell frequency observed in many inflammatory conditions. It is tempting to speculate that persistent inflammation and microbial colonization in COPD lungs could result in aberrant activation of MAIT cells leading to exhaustion and loss, as well as inappropriate recruitment of the adaptive lung immune response. Loss of MAIT cells could subsequently be an important factor in the inability to reverse tissue pathology observed in COPD lungs, due to the loss of their function in tissue repair. In this way, MAIT cells could be important early immune contributors supporting the Goldilocks hypothesis of COPD pathogenesis proposed by Curtis et al., where too strong or too weak adaptive immune response can lead to worsened symptoms of COPD^17^. Here, we considered how changes to large airway epithelial cells, the first line of defense against external assaults important to the development of COPD pathology, including cigarette smoke and microbial infection, alter MAIT cell activation.

We found that acute exposure of BEC to CS generally resulted in decreased MAIT cell responses. This finding was particularly striking in the context of microbially-infected BEC, where despite significantly increased infection of BEC exposed to cigarette smoke, we observed significantly decreased MAIT cell response. We and others have repeatedly demonstrated *in vitro* and directly *ex vivo* that increased microbial antigen or infection of healthy, untreated cells results in increased MAIT cell responses (e.g.^39^). During microbial infection, MAIT cells are thought to play an important early role in immune response; for example, through the recruitment of cells like inflammatory monocytes to the site of infection^44,45^. Delayed recruitment of adaptive immune responses in the lungs of otherwise healthy smokers and in the context of COPD exacerbations could allow for microbial persistence, inappropriately amplifying and prolonging lung inflammation.

We also observed greater microbe-independent MAIT cell responses to BEC than those observed in response to airway epithelial cell lines. These responses were also significantly higher in response to BEC from COPD lungs. We initially hypothesized this would be the result of cytokine-mediated MAIT cell activation due to reports of increased expression of cytokines like IL-18 in COPD lungs^46-48^. To our surprise, these MAIT cell responses did not require IL-12 and IL-18, but were in fact dependent on MR1. Therefore, we examined MR1 gene expression. Little is known about the regulation of MR1 gene expression, although it is known that overexpression of MR1 increases MR1-dependent MAIT cell responses (e.g. Huber et al.^40^). Additionally, genome-wide studies have identified *MR1* as a gene with altered expression or methylation status in the context of e-cigarette smoking^49^ and COPD lungs^50^. Although our sample size was not sufficiently powered for statistical significance in this area, our RT-PCR data suggests the possibility for increased *MR1* expression in BEC from COPD donors. While BEC from smoker donors did express significantly more *MR1*, we did not observe a corresponding increase in MAIT cell response. There was also no impact of acute exposure to cigarette smoke on baseline *MR1* expression in donors from any group, complicating the argument for a role of altered transcriptional regulation of *MR1* in dysregulated induction of MAIT cell IFN-γ production by uninfected BEC.

We considered other possible explanations for the increased microbe-independent, MR1-dependent responses observed in BEC from COPD lungs. One group has posited the possibility that long-term tissue damage caused by cigarette smoke could lead to the production of T cell neoantigens that contribute a potential autoimmune component to COPD-associated inflammation^51^. There has not yet been an endogenous MR1 ligand identified, however, increasing evidence from cancer MAIT cell biology suggests the existence of self ligands that can be modified in disease states^52^. Because neoantigens are already known to be important MR1 ligands^53^, the role of potential novel MR1 neoantigens produced in the context of damage from long-term cigarette smoke and COPD inflammation should be an avenue of interest. Given the small molecule nature of MR1 ligands, we initially hypothesized that cigarette smoke itself could contain novel ligands. However, absent other antigens, we did not observe any significant increase in MR1 expression of CSE-treated BECs. Furthermore, our functional data demonstrate that, if CS did contain MR1 ligands, they would not be MAIT-TCR stimulatory. If anything, exposure to cigarette smoke decreased the microbe-dependent MAIT cell responses, suggesting that any putative ligands in cigarette smoke would be antagonistic. Alternately, acute exposure to cigarette smoke resulted in an increase in 6-FP-mediated MR1 surface translocation. This increase could be mediated by CS through altered MR1 trafficking influencing ligand availability or access to putative chaperones for MR1. Together, these results demonstrate that short-term and long-term exposure to cigarette smoke could distinctly influence MR1 antigen presentation leading to dysregulated MAIT cell responses.

The mechanisms underlying COPD onset in some chronic smokers, but not others, remain unclear^17,54^. Dysfunctional MAIT cell activation could play a role in early development of COPD-associated inflammation. Absent microbial stimulus, the greater overall MAIT cell response to COPD BEC suggests that hyper-active MAIT cells could facilitate inappropriate airway inflammation, possibly through the recruitment of inflammatory monocytes and neutrophils. Conversely, the hypoactivation of MAIT cells in response to infected and CS-exposed COPD BEC could permit microbial colonization and promote chronic stimulation of innate inflammation. In the broader pulmonary context, altered immune signaling from diverse innate and adaptive cell populations (such as alveolar macrophages and neutrophils) may contribute to MAIT cell dysregulation. Our study was limited to exploring MR1 antigen presentation by primary BEC to a healthy MAIT cell clone. Future exploration of inflammatory signaling between primary MAIT and other immune cells from COPD and smoker donors may reveal further insight into COPD development. In conclusion, we demonstrate that MR1-dependent MAIT cell responses to BEC are altered in the context of COPD and cigarette smoke exposure. Understanding these impacts on MAIT cell activation may inform future therapies to treat these critically important pulmonary diseases.

## Materials and Methods

### Human subjects

This study was conducted according to the principles expressed in the Declaration of Helsinki. Study participants, protocols and consent forms were approved by Oregon Health &Science University Institutional Review Board (IRB00000186). Written and informed consent was obtained from all donors. Human participants are not directly involved in the study. Healthy adults were recruited from among employees at Oregon Health &Science University as previously described to obtain human serum^55^.

### Cells and Reagents

Primary cells were purchased commercially from Lonza Bioscience or collected from lung tissue obtained from the Pacific Northwest Transplant Bank as previously described in ^56^. Primary BEC from healthy, COPD or smoker human donors (Table 1) were grown using Bronchial Epithelial Growth Media (CC-3170) and harvested using ReagentPack Subculture reagents (CC-5034) (Lonza). BEAS-2B bronchial epithelial cells were obtained from the American Type Culture Collection (ATCC CRL-9609) and cultured in DMEM media (Gibco) supplemented with L-glutamine (25030164, Life Technologies) and 10% heat-inactivated fetal bovine serum. BEAS-2B:doxMR1-GFP cells stably expressing an MR1-GFP construct under a doxycycline-inducible promoter^39,57^ were cultured similarly to the wildtype, with doxycycline addition 16 hours prior to harvesting for analysis. The MR1-restricted T cell clone D426G11 was generated and expanded in RPMI media (Gibco) supplemented with L-glutamine and 10% heat-inactivated human serum (“RPMI-HuS”) as previously described^33,55^.

*Streptococcus pneumoniae*^29^ was cultured on tryptic soy agar plates with 5% sheep’s blood for 15 hours. Colonies were transferred to brain heart infusion broth and cultured to OD600 between 0.55-0.65 before supplementation with 20% glycerol and storage at −80°C. *Mycobacterium smegmatis* Mc^2^155 (ATCC) was grown in 7H9 broth to late log phase, before supplementation with 20% glycerol and storage at −80°C. The following antibodies were used: for ELISPOT assays: α-MR1 (26.5, Biolegend), α-IL-12p70 (MAB219100, R&D systems), α-IL-18 (D044-3, MBL International Corporation), α-IgG2A isotype (400224, Biolegend), α-human IFN-γ (7-B6-1, MabTech); for fluorescence microscopy: α-human HLA-A,B,C (W6/32, biotinylated, Biolegend), streptavidin-AlexaFluor-647 (Life Technologies); for flow cytometry: α-MR1 (26.5, conjugated to APC, Biolegend), α-human HLA-A,B,C (W6/32, conjugated to APC, Biolegend). Phytohemagglutinin PHA-L (L4144 Sigma) was suspended in RPMI-HuS. NucBlue Cell Stain ReadyProbes (ThermoFisher) and the succinimidyl ester of AlexaFluor 488 (ThermoFisher) were used per manufacturer’s protocol for microscopy. Doxycycline (Sigma) was suspended in sterile water and used at 2 μg/ml. 6-formylpterin (6-FP, Schirck’s Laboratories) was suspended in 0.01 M NaOH and used at a final concentration of 100 μM.

### Cigarette Smoke Extract preparation

Cigarette smoke extract (CSE) was prepared as in ^58^ using research grade cigarettes (1R6F, University of Kentucky Tobacco and Health Research Foundation). Briefly, smoke is collected into a polypropylene 60 ml syringe at a rate of 1 puff per minute for a total of 10 puffs per individual cigarette. Each puff consists of drawing 35 ml of smoke over 2-second duration, then slowly infusing the gas into 25 ml RPMI media over 60 seconds. CSE (pH = 7.4) or control RPMI are then sterile filtered and stored at −20°C. Freshly-thawed aliquots of RPMI (“0% CSE”) or CSE (“30% CSE”) are diluted to 30% v/v final concentration in culture medium.

### ELISPOT assay

IFN-γ ELISPOT assays were performed as previously described^59^ with following modifications: Primary BEC or BEAS-2B cells (1e5 cells/well) were used as antigen presenting cells. BEAS-2B cells were used from frozen stocks to serve as internal controls. For antibody blocking experiments, plated cells were incubated with isotype control, α-MR1, or α-IL-12 &α-IL-18 antibodies for 4 hours prior to the addition of antigen. Where indicated, BEC were incubated with RPMI-HuS containing 0% or 30% CSE for 3 hours before addition of antigen. Cells were infected with *S. pneumoniae* or *M. smegmatis*, or treated with PHA or control medium for 1 hour at 37°C. D426G11 MAIT cell clones were added (1.5e5 cells/well) before overnight incubation at 37°C. Following extensive washing with PBS-0.05% Tween 20, plates were incubated with conjugated α-human IFN-γ antibody for 2 hours before additional washing and colorimetric development. IFN-γ spot-forming units were quantified by AID ELISPOT reader.

### Fluorescence microscopy

Primary BEC were seeded directly in #1.5 glass bottom chamber slides (Nunc). Upon growth to 60-80% confluency, culture medium was replaced with pre-warmed BEGM containing 0% or 30% CSE and incubated for 3 hours at 37°C. Cells were then infected for 3 hours with AlexaFluor 488-labeled *S. pneumoniae* as previously described^29^ before washing with PBS to remove unattached bacteria and fixation with 4% paraformaldehyde for 1 hour. Slides were stained with α-HLA-A,B,C antibody and NucBlue nuclear stain, then stored at 4°C in Tris-buffered saline until imaging. Images were acquired using a high-resolution wide-field CoreDV microscope (Applied Precision) with CoolSNAP ES2 HQ (Nikon) and approximately 20 fields per condition were selected by unbiased nuclear stain. Images were taken in Z stacks in a 1024 × 1024 format using a 60 × Plan Apo N objective (NA 1.42) and an iterative algorithm was used to deconvolve the images using an optical transfer function of 10 iterations (Softworx, Applied Precision).

### Real-time quantitative PCR (RT-PCR)

RNA isolation was performed with the RNeasy Plus kit (Qiagen) and cDNA synthesis was completed with the High Capacity cDNA Reverse Transcription Kit (Life Technologies) following the manufacturers’ protocols. Real-time PCR was performed using Taqman gene expression assays for *MR1* (Hs01042278_m1), obtained from Applied Biosystems. Reactions were run in triplicate. Expression data were normalized to *HPRT1* (Hs02800695_m1) and relative expression levels for each target gene were determined using the 2^-ΔΔCt^ method^60^.

### Surface MR1 and MHC-Ia flow cytometry

Primary BEC, WT BEAS-2B cells, and BEAS-2B:doxMR1-GFP cells were plated in 6-well tissue culture plates. Where indicated, cell medium was replaced with pre-warmed medium containing 0% or 30% CSE for 3 hours at 37°C prior to overnight incubation with 100 uM 6-FP. After 16 hours incubation, cells were harvested and suspended in FACS buffer containing 2% heat-inactivated human serum, 2% heat-inactivated goat serum, and 0.5% heat-inactivated FBS for 30 minutes on ice. Samples were then stained with APC-conjugated 26.5 α-MR1 antibody, α-W6/32 antibody, or isotype control antibody for 40 minutes at 4°C. Cells were washed, fixed with 1% paraformaldehyde and analyzed with a Beckman Coulter CytoflexS. All analyses were performed using FlowJo10 (TreeStar).

### IL-18 expression

IL-18 immunoassay was preformed using the ProQuantum Human IL-18 Immunoassay Kit (A35613, Invitrogen) per manufacturer’s protocols with a 1:3 dilution of supernatants collected from primary BEC following indicated infection with *S. pneumoniae* and incubation with D426G11 MAIT cell clones.

### Data analysis

ELISPOT statistical analysis was performed using R 3.6.3 and packages such as ggplot2, dplyr, lem4, afex, emmeans and multcome. The lmer function in lme4 package was used to do the analysis and first the best fitting model structure was searched using the anova function implementing likelihood ratio test. The linear mixed effects model with square root transformation of SFU was used to analyze the data. Post-hoc tests to determine group differences were run using function glht from package multcomp (to compare groups and perform a z-test) and function emmeans (Estimated Marginal Means) from package emmeans (to perform t-tests for pairwise comparisons). All other data were analyzed using Prism 8 (GraphPad) or R 4.0 using packages such as ggplot2, kableExtra, ggh4x, and ggbeeswarm. Statistical significance was determined as indicated by pairwise t test or two-way ANOVA with Bonferroni’s multiple comparisons, using α=0.05. All images were analyzed using Imaris (Bitplane) as in ^39,40^.

## Acknowledgements

The D426G11 MAIT cell clone was a kind gift from Dr. David Lewinsohn. We thank Dr. Suil Kim at VA Portland Health Care System for technical advice on cigarette smoke extract preparation and use. We also thank Dr. Yalda Zarnegarnia and Jack Wiedrick of the Oregon Health &Science University Biostatistics &Design Program for their assistance in statistical analysis. The DeltaVision Core DV microscope is supported in part by NIH S10 RR023432.

## Funding

This work was supported by VA Merit Award I01 CX001562 (MJH), and in part by NIH R01 AI129976 (MJH) and NIH T32 1T32GM137794-05 (MEH). The contents do not represent the views of the U.S. Department of Veterans Affairs or the U.S. government.

---

**Supplemental Figure 1:**
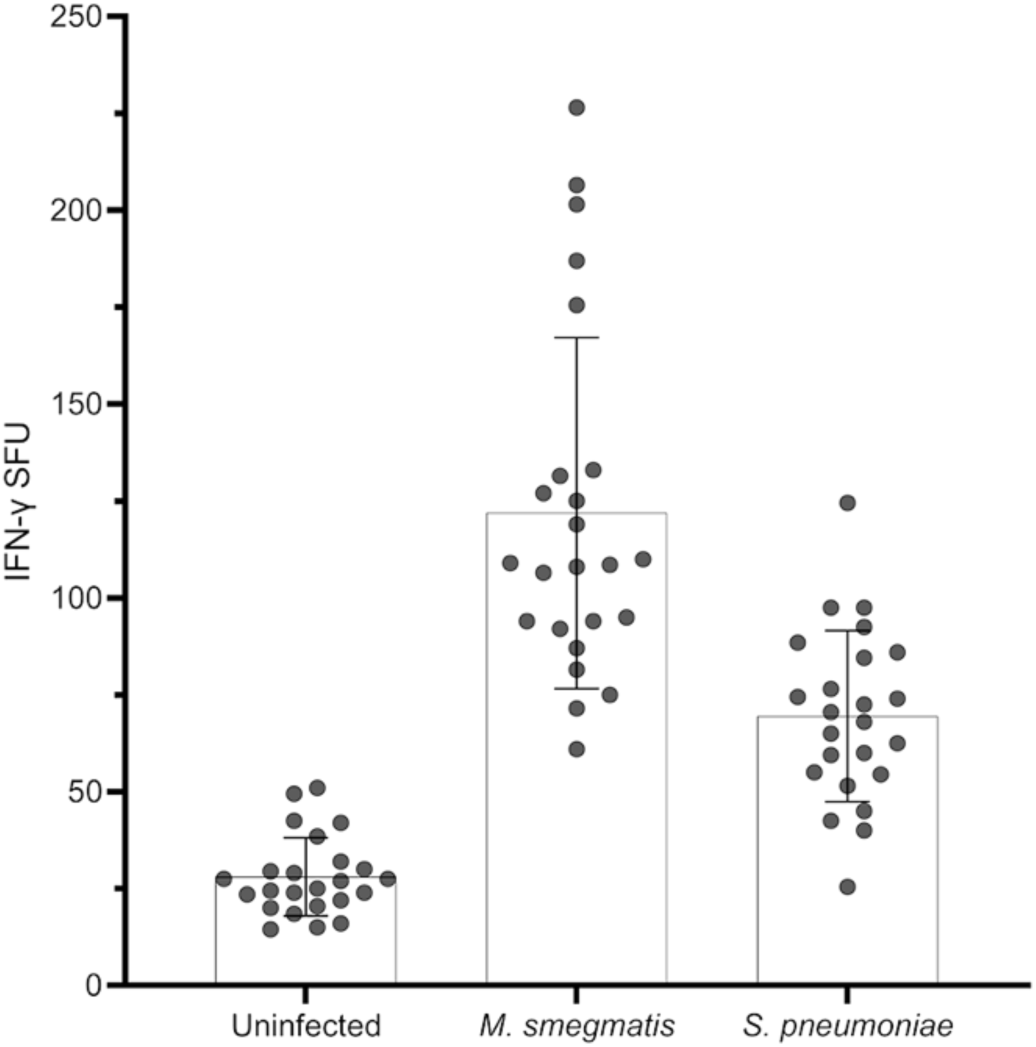
BEAS-2b cells elicit MAIT cell responses. BEAS-2b cells were incubated with media control (“Uninfected”) or infected with *M. smegmatis* (0.05μl/well) or *S. pneumoniae* (20 MOI) for one hour prior to addition of MAIT cells in an IFN-γ ELISPOT assay. Data points are the mean IFN-γ spot-forming units (SFU) of two technical replicates each from 24 experiments.

**Supplemental Figure 2:**
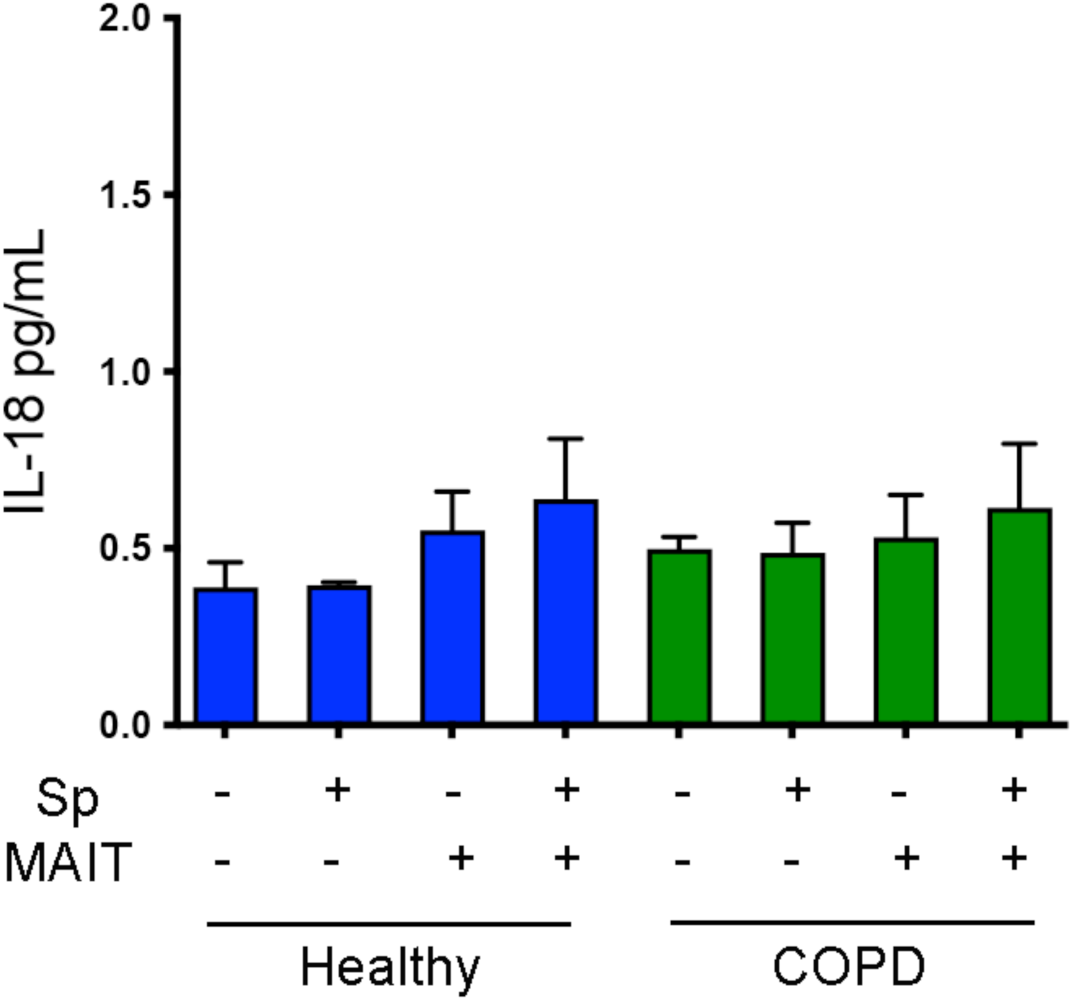
IL-18 secretion by primary BEC. Cell supernatants were collected from a representative healthy, COPD, and smoker donor BEC treated as indicated. IL-18 secretion was quantified from cell supernatants by immunoassay.

**Supplemental Figure 3:**
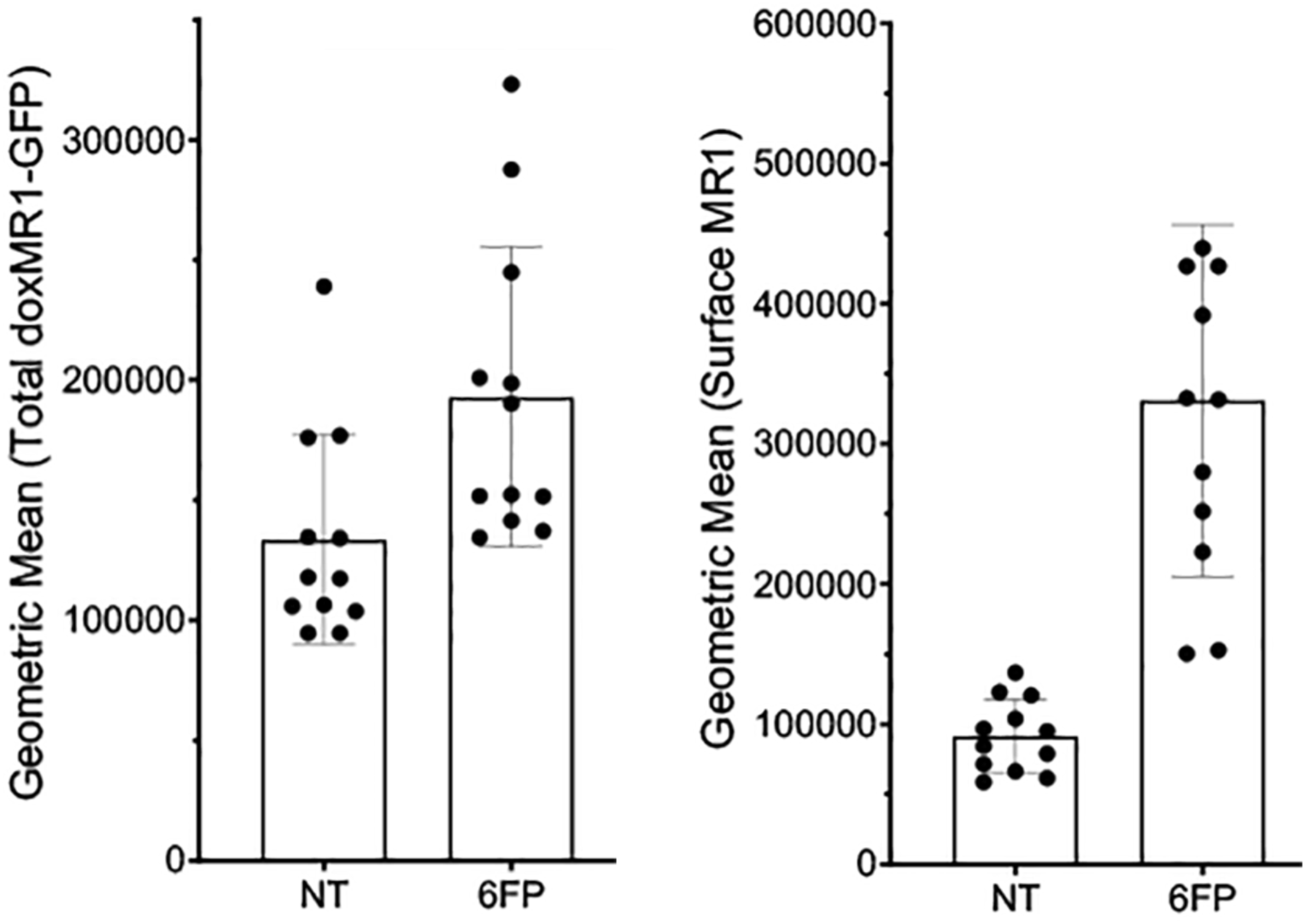
Expression of MR1 and MHC-Ia in the BEAS-2B cell line. BEAS-2B:doxMR1-GFP cells were treated with doxycycline 24 hours prior to overnight incubation with 6-FP. Cells were stained and analyzed by flow for total cellular doxMR1-GFP expression (left) or for surface expression of MR1 (right).

## Notes

### Competing Interest Statement

The authors have declared no competing interest.

